# The development of gut microbiota in ostriches and its association with juvenile growth

**DOI:** 10.1101/270017

**Authors:** Elin Videvall, Se Jin Song, Hanna M. Bensch, Maria Strandh, Anel Engelbrecht, Naomi Serfontein, Olof Hellgren, Adriaan Olivier, Schalk Cloete, Rob Knight, Charlie K. Cornwallis

**Affiliations:** Department of Biology, Lund University, Lund, Sweden; Department of Pediatrics, University of California San Diego, La Jolla, CA, USA; Directorate Animal Sciences, Western Cape Department of Agriculture, Elsenburg, South Africa; Western Cape Agricultural Research Trust, Elsenburg, South Africa; Klein Karoo International, Research and Development, Oudtshoorn, South Africa; Department of Animal Sciences, Stellenbosch University, Matieland, South Africa; Department of Computer Science & Engineering, University of California San Diego, La Jolla, CA, USA; Center for Microbiome Innovation, University of California San Diego, La Jolla, CA, USA

**Keywords:** gut microbiome, struthio camelus, development, succession, colonisation, weight

## Abstract

The development of gut microbiota during ontogeny in vertebrates is emerging as an important process influencing physiology, immune system, health, and adult fitness. However, we have little knowledge of how the gut microbiome is colonised and develops in non-model organisms, and to what extent microbial diversity and specific taxa influence changes in fitness-related traits. Here, we used 16S rRNA gene sequencing to describe the successional development of the faecal microbiota in juvenile ostriches (*Struthio camelus*; n = 71) over their first three months of life, during which time a five-fold difference in weight was observed. We found a gradual increase in microbial diversity with age, an overall convergence in community composition among individuals, multiple colonisation and extinction events, and major taxonomic shifts coinciding with the cessation of yolk absorption. In addition, we discovered significant but complex associations between juvenile growth and microbial diversity, and identified distinct bacterial groups that had positive (Bacteroidaceae) and negative (Enterobacteriaceae, Enterococcaceae, Lactobacillaceae) correlations with the growth of individuals at specific ages. These results have broad implications for our understanding of the development of gut microbiota and its association with juvenile growth.

## Introduction

The gastrointestinal tract of vertebrates is considered to be largely sterile at the time of birth (Perez-Muñoz *et al.* 2017; cf. Jiménez *et al.* 2008) and subsequently colonised by a wide array of micro-organisms, collectively termed ‘the gut microbiota’. The gut microbial composition during early life has been shown to have major influences on the health and phenotype of adults through its effects on gut morphology, metabolism, immune system development, and brain development (Dominguez-Bello *et al.* 2010; Heijtz *et al.* 2011; Russell *et al.* 2012; Cho *et al.* 2012; Cox *et al.* 2014). For example, animals prevented from acquiring gut bacteria suffer from smaller intestines with thinner gut walls, smaller lymph nodes, a poorly developed immune system, and reduced organ sizes including heart, lungs, and liver (Gordon & Pesti 1971; Mitsuhiro & Jun-ichi 1994; Macpherson & Harris 2004). Similarly, animals with a poorly developed gut microbiota have an altered metabolism (Cox *et al.* 2014) and are more susceptible to infection by pathogenic bacteria, viruses, and eukaryotes (Sprinz *et al.* 1961; Inagaki *et al.* 1996; Round & Mazmanian 2009). Given the crucial effects of gut bacteria on hosts, it is important to characterise how, when, and by what microbes the gut is colonised, to examine whether variation in this process explains differences in host development.

The majority of research on the microbial colonisation of the gut during host development, and its associated effects on fitness, has been on humans, as well as domesticated and model laboratory animals. In some animals it has been found that the diversity of the gut microbiota increases with age during ontogeny, whereas in others the reverse is true. For example, in mice and humans, colonisation is initiated during birth, where the mother’s vaginal and skin microbiota are important sources of bacteria (Sommer & Bäckhed 2013; Pantoja-Feliciano *et al.* 2013; Kundu *et al.* 2017). Seeding of microbes continues through lactation, and during the first year of life the human gut microbiome remains relatively simple with low diversity, and varies markedly across individuals and over time. The gut microbiome subsequently shifts during weaning towards an adult-like bacterial community and becomes more stable (Sekirov *et al.* 2010; Koenig *et al.* 2011; Yatsunenko *et al.* 2012). In contrast, in species such as zebrafish (*Danio rerio*) and African turquoise killifish (*Nothobranchius furzeri*), the alpha diversity and richness of the gut microbiota is highest in neonatal juveniles and subsequently decreases during maturation (Stephens *et al.* 2016; Smith *et al.* 2017). Similar to fish, juvenile birds rely heavily on environmental and dietary sources for acquiring the initial gut microbes (Lu *et al.* 2003; Yin *et al.* 2010). However, depending on the level of parental care, some bird species may receive significant microbial contributions from their parents, via for example regurgitation or shared nest environment (Godoy-Vitorino *et al.* 2010; van Dongen *et al.* 2013; Dewar *et al.* 2017).

Differences in the colonisation of gut microbiota during development have been shown to have pronounced and highly variable long-term effects on hosts. It has been found that bacterial diversity in the gut promote host development and growth by enabling greater resource acquisition and preventing domination by certain bacteria (Ley *et al.* 2006; Lozupone *et al.* 2012; Foster *et al.* 2017). For example, studies have demonstrated that germ-free animals require a higher calorific intake to attain the same growth as hosts with a normal microbial diversity (Wostmann *et al.* 1983; Bäckhed *et al.* 2004; Shin *et al.* 2011; Sommer & Bäckhed 2013). Conversely, it has been suggested that a reduced diversity in the gut microbiota may increase growth and accelerate host development. This idea is supported by numerous studies from the agricultural industry where higher growth rates have been achieved in farm animals by eliminating gut bacteria with antibiotics, a common practice since the 1950s (Gaskins *et al.* 2002; Dibner & Richards 2005; Lin *et al.* 2013). Likewise, supplementing wild animals with antibiotics has also been associated with positive effects on growth (Potti *et al.* 2002; Kohl *et al.* 2017). Pinpointing the exact mechanisms through which gut microbes influence host growth has, however, been problematic as antibiotics in some cases increase microbial diversity (Crisol-Martínez *et al.* 2017; Kohl *et al.* 2017). Similarly, some probiotic supplements have led to an increase in animal growth while others are associated with a reduction in growth (Million *et al.* 2012; Angelakis *et al.* 2013). These alternative predictions and conflicting reports on gut microbiota and animal growth highlight the need for separating the effects of diversity and specific bacterial groups. For example, different taxa may be associated with either an increase or decrease in host metabolism and/or intestinal immune responses, which may engender highly different effects on juvenile development and growth.

In this study, we evaluated the developing gut microbiome of ostrich (*Struthio camelus*) chicks over time and in relation to their growth. We performed repeated faecal sampling of individually tagged ostriches in a research rearing facility from their first week after hatching until 12 weeks of age, which constitutes the critical developmental phase in this species (Verwoerd *et al.* 1999; Cloete *et al.* 2001). Ostriches are the largest living bird species and, together with other paleognaths, are basal in the phylogeny of birds. They are a valuable economic resource being farmed for feathers, meat, eggs, and leather, yet have only been kept in captivity for a short period of time relative to other agricultural animals (Cloete *et al.* 2012). Their chicks are highly precocial, allowing them to be raised independently from parents, and they reach sexual maturity from two years of age. Ostriches also have one of the largest variations in offspring growth rate across birds, even in controlled environments (Deeming & Ayres 1994; Skadhauge & Dawson 1999; Bonato *et al.* 2009; Engelbrecht *et al.* 2011), and are known to suffer from bacterial gut infections (Verwoerd 2000; Keokilwe *et al.* 2015). These traits make the ostrich an excellent organism for investigating host-microbiota associations, including the effects of gut microbiota on juvenile growth and development.

## Materials and methods

### Experimental setup

Juvenile ostriches were kept under controlled conditions at the Western Cape Department of Agriculture’s ostrich research facility in Oudtshoorn, South Africa. Chicks were obtained from a batch of artificially incubated eggs that hatched on Sep 30^th^ 2014. A total of 234 individuals were monitored from hatching date until three months of age (12 weeks) in four groups that contained around 58 chicks each at the start of the experiment. The groups were kept in indoor pens of approximately 4×8 m with access to outdoor enclosures with soil substrate during the day. To reduce potential environmental variation on the development of the gut microbiota, all individuals were reared under standardized conditions with *adlibitum* access to food and fresh water during the daytime. The chicks received a standardised pelleted pre-starter ration, and the adult birds were given pelleted breeder ration and were kept in a different area separate from the chick facility. All procedures were approved by the Departmental Ethics Committee for Research on Animals (DECRA) of the Western Cape Department of Agriculture, reference no. R13/90.

### Sample collection

Faecal samples in this study were collected from chicks during the following ages: week 1, 2, 4, 6, 8, and 12. In addition, we sampled fresh faeces from five adult individuals kept in large outside enclosures. The sex and age of the adults are not known, but the samples were collected from sexually mature, breeding individuals. All faecal samples were collected in empty plastic 2 ml micro tubes (Sarstedt, cat no. 72.693) and stored at −20 °C within two hours of collection. They were subsequently transported on ice to a laboratory and stored at −20 °C. Detailed sample collection has been described by Videvall *et al.* (2017a). Weight measurements of the ostriches were retrieved during each sampling event. At the final time point (week 12), the smallest ostrich chick weighed 6 kg while the largest weighed 30 kg, representing a five-fold difference in body mass (mean = 18 kg).

Throughout the course of the trial, a large number of individuals died (n = 72) primarily from suspected disease, as is common in ostrich rearing facilities (Verwoerd *et al.* 1999; Cloete *et al.* 2001). In addition, 10 chicks were randomly selected for euthanization and dissection at 2, 4, 6, 8, 10, and 12 weeks of age (n = 60), to act as age-matched controls for the diseased individuals. The gut samples of the diseased and euthanized juveniles have been analysed in a different study (Videvall *et al.* unpublished data). To investigate the development and maturation of ostrich microbiomes, the samples used in this study were those retrieved from euthanized (control) individuals and from individuals that survived the entire period (n = 71). The faecal samples from the individuals that died from suspected disease were not included.

### DNA isolation, library preparation, and amplicon sequencing

We prepared sample slurries for all sample types with guidance from Flores *et al.* (2012) and subsequently extracted DNA using the PowerSoil-htp 96 well soil DNA isolation kit (Mo Bio Lboratories, cat no. 12955-4) as recommended by the Earth Microbiome Project (www.earthmicrobiome.org). For full details please see Videvall *et al.* (2017a). Amplicon libraries for sequencing of the 16S rRNA V3 and V4 regions were prepared using Illumina fusion primers containing the target-specific primers Bakt_341F and Bakt_805R (Herlemann *et al.* 2011) according to the Illumina 16S Metagenomic Sequencing Library Preparation Guide (Part # 15044223 Rev.B). The samples were sequenced as 300 bp paired-end reads over three sequencing runs on an Illumina MiSeq platform at the DNA Sequencing Facility, Department of Biology, Lund University, Sweden. A total of 277 faecal samples plus 4 negative samples were part of this study (Table S1).

### Data processing

The 16S amplicon sequences were quality-screened using FastQC (v. 0.11.5) (Andrews 2010) together with MultiQC (Ewels *et al.* 2016). Primers were removed from the sequences using Trimmomatic (v. 0.35) (Bolger *et al.* 2014) and the forward reads were retained for analyses. Quality filtering of the reads was executed using the script multiple_split_libraries_fastq.py in QIIME (v. 1.9.1) (Caporaso *et al.* 2010). All bases with a Phred score < 25 at the 3’ end of reads were trimmed and samples were multiplexed into a single high-quality multi-fasta file.

Operational taxonomic units (OTUs) were assigned and clustered using Deblur (v. 1.0.0) (Amir *et al.* 2017). Deblur circumvents the problems surrounding clustering of OTUs at an arbitrarily threshold by obtaining single-nucleotide resolution OTUs (100% sequence identity approach) after correcting for Illumina sequencing errors. This results in exact sequence variants (ESVs), also called amplicon sequence variants (ASVs), oligotypes, zero-radius OTUs (ZOTUs), and sub-OTU (sOTUs). In order to avoid confusion, we call these units OTUs, but note that they differ from the traditional 97% clustering approach as they provide more accurate estimates (Edgar 2017; Amir *et al.* 2017; Callahan *et al.* 2017). The minimum reads-option was set to 0 to disable filtering inside Deblur, and all sequences were trimmed to 220 bp. We used the biom table produced after both positive and negative filtering, which by default removes any reads containing PhiX or adapter sequences, and only retains sequences matching known 16S gene sequences. Additionally, PCR-originating chimeras were filtered inside Deblur (Amir *et al.* 2017).

Taxonomic assignment of OTUs was performed using the Greengenes database (v. 13_8) (McDonald *et al.* 2012). We removed the following OTUs from all samples: all OTUs present in the negative (blank) samples (n = 95), all OTUs classifying as mitochondria (n = 7), all OTUs classifying as chloroplast (n = 18), all OTUs that only appeared in one sample, and finally all OTUs with a total sequence count of less than 10. These filtering steps removed in total approximately 47,000 OTUs, with 4,338 remaining for analyses. All samples were retained since none exhibited low read coverage (the lowest coverage obtained was 1,799 reads after filtering). Further, the sequence coverage per sample (mean number of filtered reads = 15,480) showed no differences across ages (ANOVA: F = 2.01, p = 0.064). Analyses were evaluated with both rarefied and non-rarefied data, which produced extremely similar and comparable results. We therefore present the results from the non-rarefied data in this study.

### Data analyses

All analyses were performed in R (v. 3.3.2) (R Core Team 2017). We calculated OTU richness (observed OTUs) and alpha diversity (Shannon index) using absolute abundance of reads, and distance measures with Bray-Curtis and weighted UniFrac methods (Bray & Curtis 1957; Lozupone & Knight 2005) on relative read abundances in phyloseq (v. 1.19.1) (McMurdie & Holmes 2013). Differences between the microbiomes of juvenile samples were examined using permutational multivariate analysis of variances (PERMANOVA) on weighted UniFrac distances using the adonis function in vegan (v. 2.4-2) (Oksanen *et al.* 2017) with 1000 permutations. Effects of age on the microbiome were evaluated by Z-transforming age in weeks and fitting a linear and a quadratic age term. Differences in dispersion between age groups were tested with the multivariate homogeneity of group dispersions test (betadisper) in vegan (Oksanen *et al.* 2017), followed by the Tukey’s ‘Honest Significant Difference’ method. Distances in microbiomes across individuals within and between age groups were calculated using Bray-Curtis metrics on relative abundances. All comparisons between samples from the same individual were excluded prior to calculating distance metrics.

To evaluate bacterial abundances, we first modelled counts with a local dispersion model and normalised per sample using the geometric mean (according to the DESeq2 manual) (Love *et al.* 2014). Differential abundances between juvenile age groups were subsequently tested in DESeq2 with a negative binomial Wald test, while controlling for individual ID of birds, and with the beta prior set to false (Love *et al.* 2014). The results for specific comparisons were extracted (e.g. week 1 versus week 2) and p-values were corrected with the Benjamini and Hochberg false discovery rate for multiple testing (Benjamini & Hochberg 1995). OTUs were labelled significantly differentially abundant if they had a corrected p-value (q-value) < 0.01. The test between week 12 juveniles and adults was performed without individual ID in the model as this comparison did not include any repeated data measures.

Interactions between juvenile growth (weight over time) and alpha diversity (Shannon index), and juvenile growth and bacterial richness were tested using a linear mixed-effect model from the nlme package (Pinheiro *et al.* 2016). The random effects included in the mixed models were random intercepts, random slope across age and their covariance. We further investigated the relationship between alpha diversity and weight by examining correlation coefficients between alpha diversity and weight for each age group calculated with Pearson’s correlation. To investigate the relationship between specific OTUs and weight, log-transformed sums of normalised OTU abundances of taxa (filtered on a total abundance of minimum 1000 counts) were correlated with juvenile weight by using Pearson’s correlation. The resulting p-values were corrected to q-values with the false discovery rate and taxa with q < 0.05 were considered statistically significant. Plots were made using ggplot2 (Wickham 2009).

## Results

### Age has a major influence on the gut microbiota composition of juvenile ostriches

Unsupervised Non-metric Multi-Dimensional Scaling (NMDS) and Principal Coordinates Analysis (PCoA) of Bray-Curtis and weighted UniFrac distances showed that age constitutes a major part of the variation observed in the juvenile ostrich faecal microbiome (Figure 1, Figure S1). The samples followed each other in a chronological order along the first NMDS axis, where each age group approached the microbiome of adult individuals (Figure 1A). The microbiota of individuals at week 1 showed the largest differences to all other ages on the PCoA (Figure 1B) and clustered separately from the microbiota of individuals at week 2, which in turn clustered separately from those at week 4 and subsequent weeks (Figure 1A). Samples from week 4 of age showed a slightly larger variation in spread along the second NMDS axis (Figure 1A) and the samples from the oldest ages (weeks 6, 8, and 12) showed the least differences to each other, especially when analysed with weighted UniFrac distances (Figure 1B).

**Figure 1.**
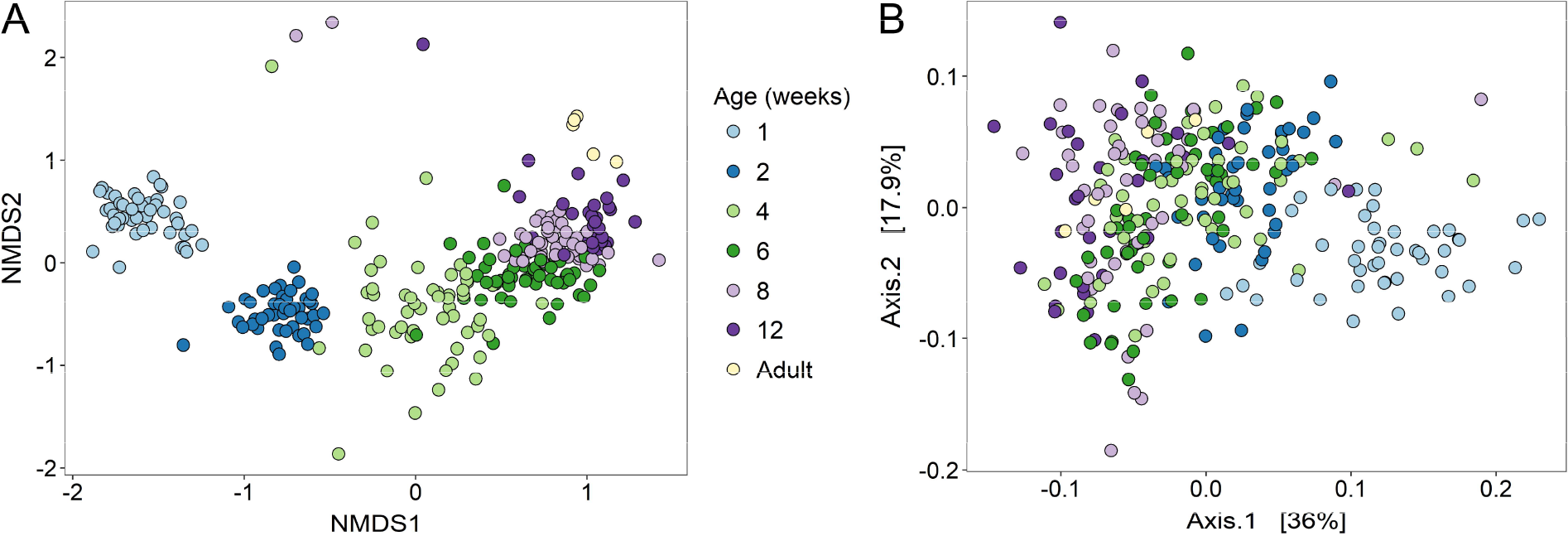
Gut microbiomes show major differences with age of hosts. (**A**) NMDS of Bray-Curtis distances and (**B**) PCoA of weighted UniFrac distances between samples. Colours indicate age of individuals in weeks and brackets in the PCoA display the percent of variance explained by the first two dimensions.

We analysed changes in the microbiome with age using a PERMANOVA on weighted UniFrac distances and found that there was a significant linear effect showing an increase in microbiome distance with age (R^2^ = 0.17; p < 0.001) and a quadratic effect (R^2^ = 0.06; p < 0.001) which indicated that changes slowed down over time. As a result, the differences between individuals increased with differences in age, where the microbiota of juveniles at week 1 was most similar to those at week 2, week 2 was most similar to those at week 1 and 4, etc.; yet the similarities between the older ages (weeks 6, 8, 12) were higher than the similarities between earlier weeks (Figure 2, Figure S2). The microbiomes within age groups were always more similar to each other than they were to all other ages, and interestingly, the youngest age, week 1, showed the most within-group similarities compared to all other ages (Figure 2, Figure S2). The degree of variation in the microbiome among individuals was similar across ages (multivariate homogeneity test of group dispersions; adjusted p = 0.203-0.999), apart from the specific comparison between week 2 and 4, which showed that week 4 was significantly more variable (adjusted p = 0.009). In line with this, our PERMANOVA showed high but non-significant variation between individuals in their microbiota (R^2^ = 0.19, p = 0.243). In contrast, sex (R^2^ = 0.003; p = 0.396) and group (R^2^ = 0.005; p = 0.081) did not have any effects on microbiota composition.

**Figure 2.**
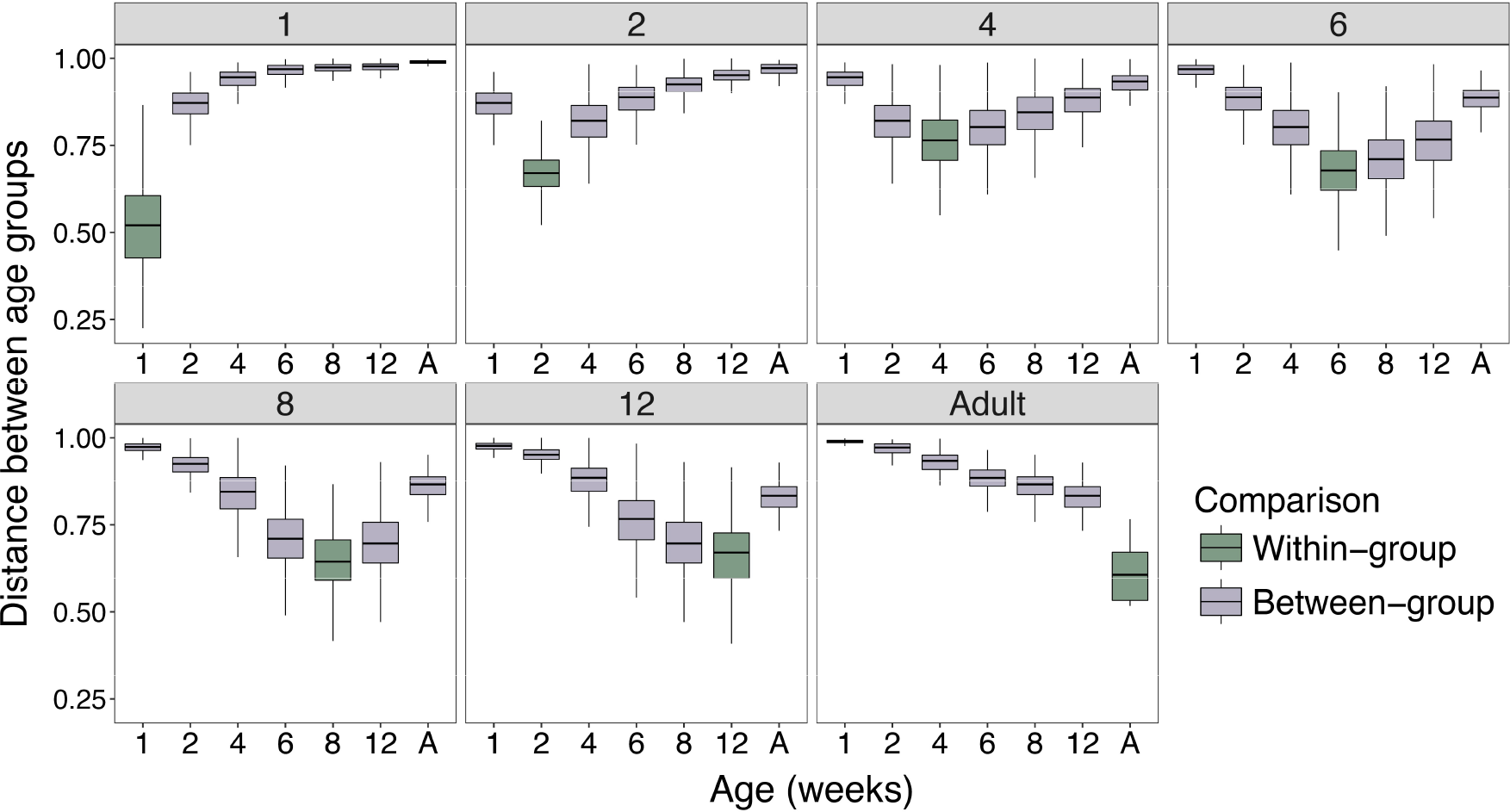
Beta diversity (Bray-Curtis distances) of gut microbiota shows most similarities within age groups. The headers show age in weeks and the x-axes show all the age comparisons, with A = Adults. Within-age group comparisons are highlighted in green (e.g. distances between all individuals at week 1) and higher values signify more dissimilar microbiomes.

Similarly to the distance measures of microbial composition, we found that the richness and alpha diversity of the gut microbiomes showed a clear and steady increase over time as individuals became older (Figure 3). The differences in alpha diversity between ages were highly significant (linear-mixed effect model (lme): age, parameter estimate (β) se = 0.08 ± 0.01, F_1, 205_ = 54.66, p < 0.0001), even after controlling for weight (lme with weight as a covariate: age, parameter estimate (β) se = 0.13 ± 0.02, F_1, 204_ = 55.35, p < 0.0001), with the samples from the earliest time point (week 1) exhibiting the lowest alpha diversity, and the adult samples the highest alpha diversity (Figure 3). Analyses of bacterial richness showed very similar results, with richness increasing with age both before (lme: age, parameter estimate (β) se = 26.89 ± 2.06, F_1, 205_ = 170.40, p < 0.0001) and after controlling for changes in weight (lme with weight as a covariate: age, parameter estimate (β) se = 35.62 ± 4.27, F_1, 204_ = 182.52, p < 0.0001).

**Figure 3.**
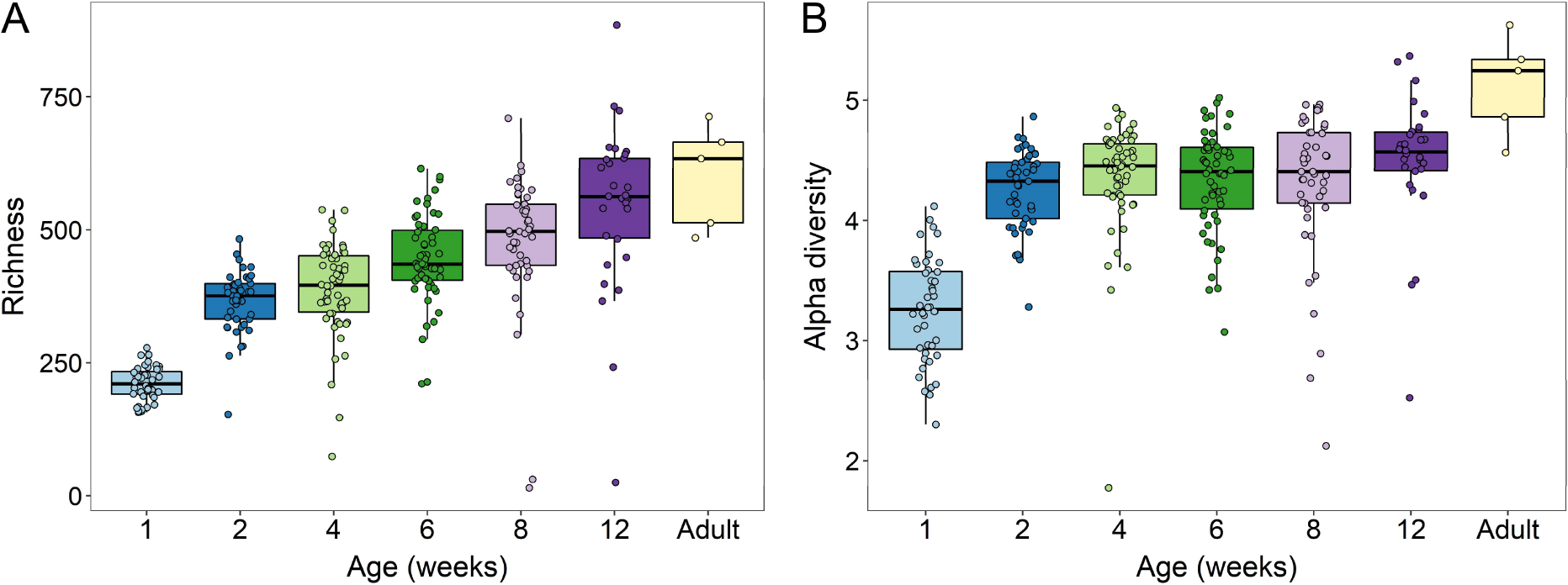
Step-wise increase of microbial diversity with host age. (**A**) Richness (observed OTUs), and (**B**) alpha diversity (Shannon index) sampled at different ages.

Investigating the taxon composition of the ostrich gut microbiome over time showed large shifts in particular for the early ages, with differences evident even at higher-levels of the taxonomy (Figure 4). Juveniles 1-week-old had high abundances of Verrucomicrobiae and Erysipelotrichi, but by week 2 these classes were already highly reduced relative to other bacteria (Figure 4). Furthermore, Planctomycetia, Verrucomicrobiae, and Gammaproteobacteria were practically absent in adults compared to juveniles (Figure 5). In contrast, Bacilli and Planctomycetia started colonising the gut around four weeks of age, after which time the relative abundances of bacterial classes remained relatively consistent with age, with Clostridia being the dominant class (Figures 4 and 5). Other taxa had more complicated relationships with age, such as the Bacteroidia, that peaked in abundance during week 2 (relative to the other classes), and subsequently decreased with age, but were higher in the adults (Figure 5).

**Figure 4.**
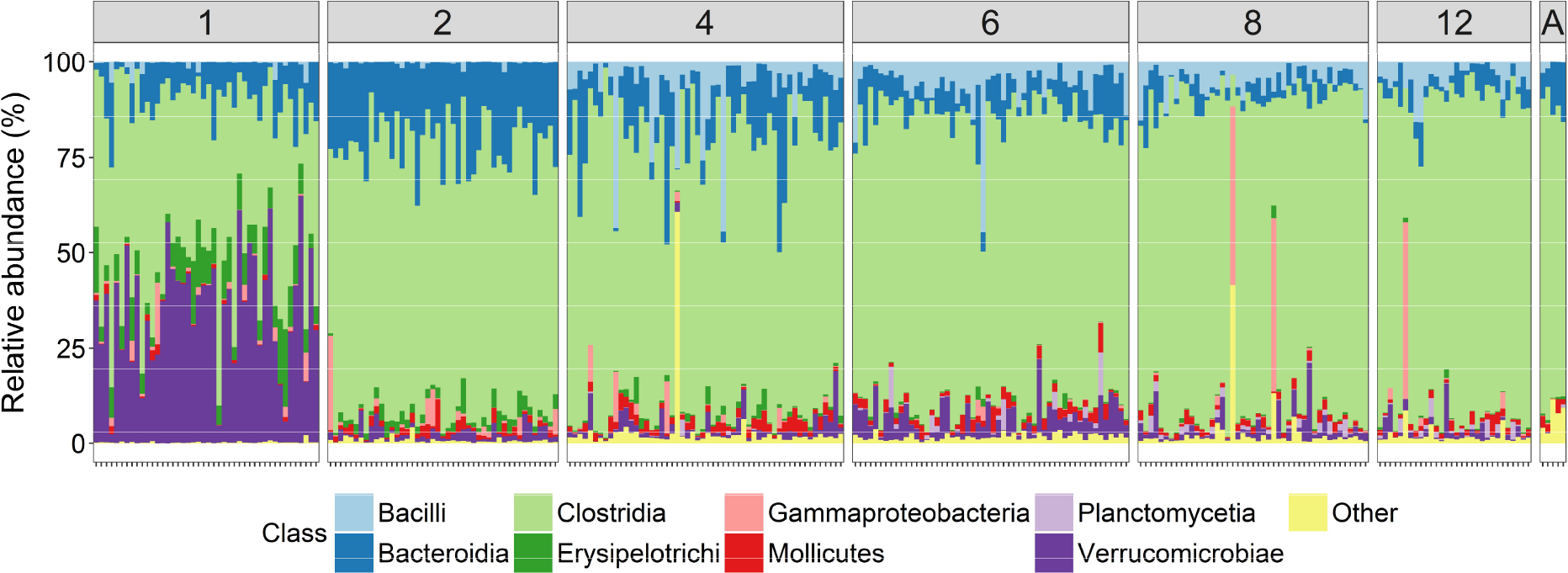
Relative abundances of bacterial groups fluctuate and gradually stabilise with increasing age of hosts. Colours represent bacterial classes and each bar displays the bacterial composition for one host individual. The headers show age in weeks, except A = Adult individuals.

**Figure 5.**
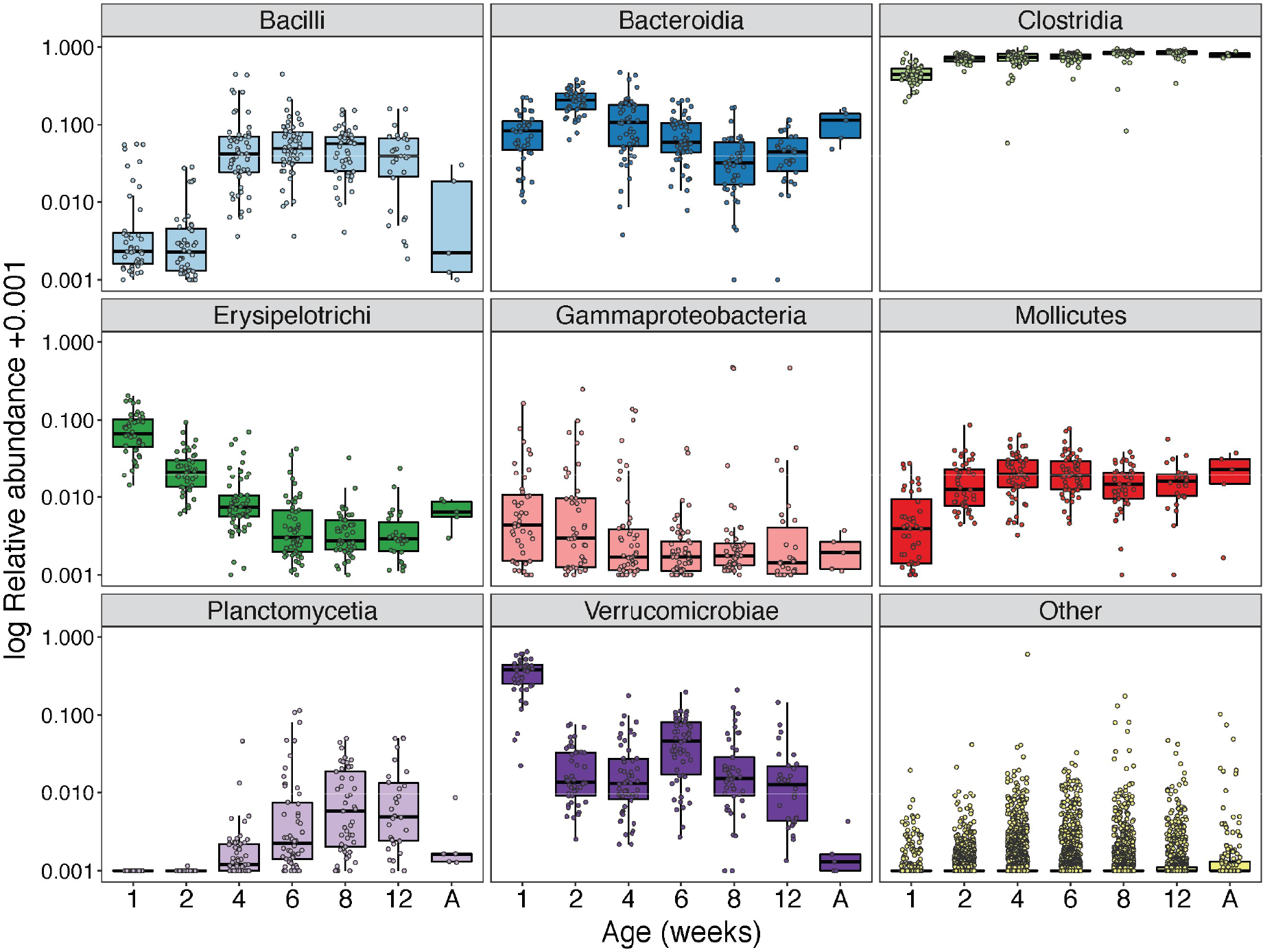
Relative abundances of different bacterial classes (log-transformed +0.001) display different trends with increasing host age. The x-axes show age in weeks, except A = Adult individuals.

Examining differences in OTU abundance between age groups produced a more detailed picture of the bacterial shifts during the development of the ostrich gut microbiome. The top most prevalent OTUs at the different ages belonged to *Akkermansia muciniphila* during week 1 (27.2%), Bacteroidales sp. during week 2 and 4 (7.1-11.9%), Clostridiaceae sp. during week 6, 8, and 12 (7.9-8.6%), and Ruminococcaceae sp. in the adults (5.1%) (Table S2). The abundance of all OTUs became more similar over time as individuals aged (Figure 6). For instance, the comparison between week 1 and week 2 showed the fewest similarities in overall OTU abundances, despite having the shortest interval between sampling events, while the comparison between week 8 and week 12 showed large similarities (Figure 6). The comparison between week 12 juveniles and adult birds displayed a large proportion of highly abundant OTUs unique to each age group (i.e. only present in either adult birds or in 12-week-old juveniles) (−1 values in Figure 6).

Negative binomial Wald tests of normalised OTU abundances between age groups closest in time resulted in a large number of significant differentially abundant OTUs. Specifically, several OTUs were more abundant in 2-week-old juveniles compared to 1-week-olds (Figure 7; Table S3), with the most significant OTUs coming from the families Ruminococcaceae and Christensenellaceae, and 16 OTUs within Bacteroidia (families Bacteroidaceae, S24-7, Rikenellaceae, and Odoribacteraceae) were significantly more abundant at week 2 relative to week 1 (Table S3). The analysis between week 2 and week 4 yielded a large number of significant OTUs (n = 498), of which the majority (70.7%) were again more abundant in the older age group, demonstrating microbial establishment. Notably, almost half (47.4%) of the significant OTUs were completely absent at week 2 but were present at week 4, including, for example, OTUs within Actinobacteria and Planctomycetia (Table S4). At week 6 there were again numerous colonisations (n = 166), mostly within the classes Clostridia and Mollicutes, but some OTUs had gone locally extinct (n = 68) (Table S5). By week 8, extinction (n = 88) and colonisation events (n = 80; Table S6) were approximately equal, and by week 12 changes in OTU abundance had slowed down with fewer significant differentially abundant OTUs relative to week 8 (n = 182; Table S7). The final comparison between week 12 juveniles and adults yielded 60 significant OTUs, of which all except one (*Aerococcus* sp.) were practically absent in adults (Figure 7; Table S8), potentially resulting from the smaller number of adult samples. As a specific example of the pattern of colonisation in juveniles, two archaea *Methanocorpusculum* OTUs were completely absent in younger individuals but appeared at week 6 and 8, respectively, whereas one archaea *Methanobrevibacter* OTU appeared at week 2 and increased markedly in abundance over time.

**Figure 6.**
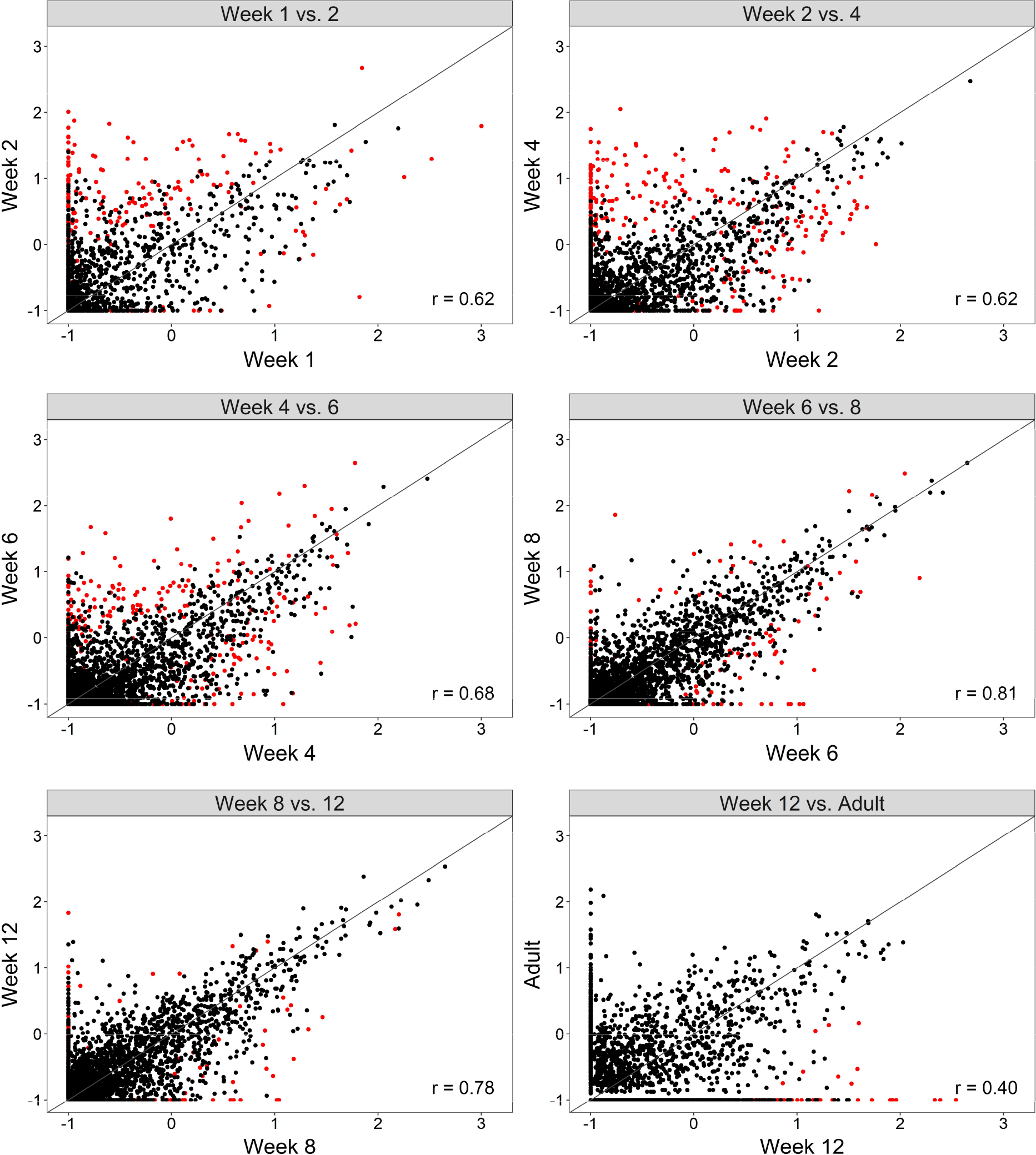
Abundances of all OTUs increase in similarity with increasing host age. Axes show log-transformed mean normalised OTU abundances (+0.1) between age groups closest in time. Significant differentially abundant OTUs are highlighted in red and the 1:1 relationship is indicated by the diagonal line. The Pearson’s correlation coefficients for these values are provided for each comparison (r).

**Figure 7.**
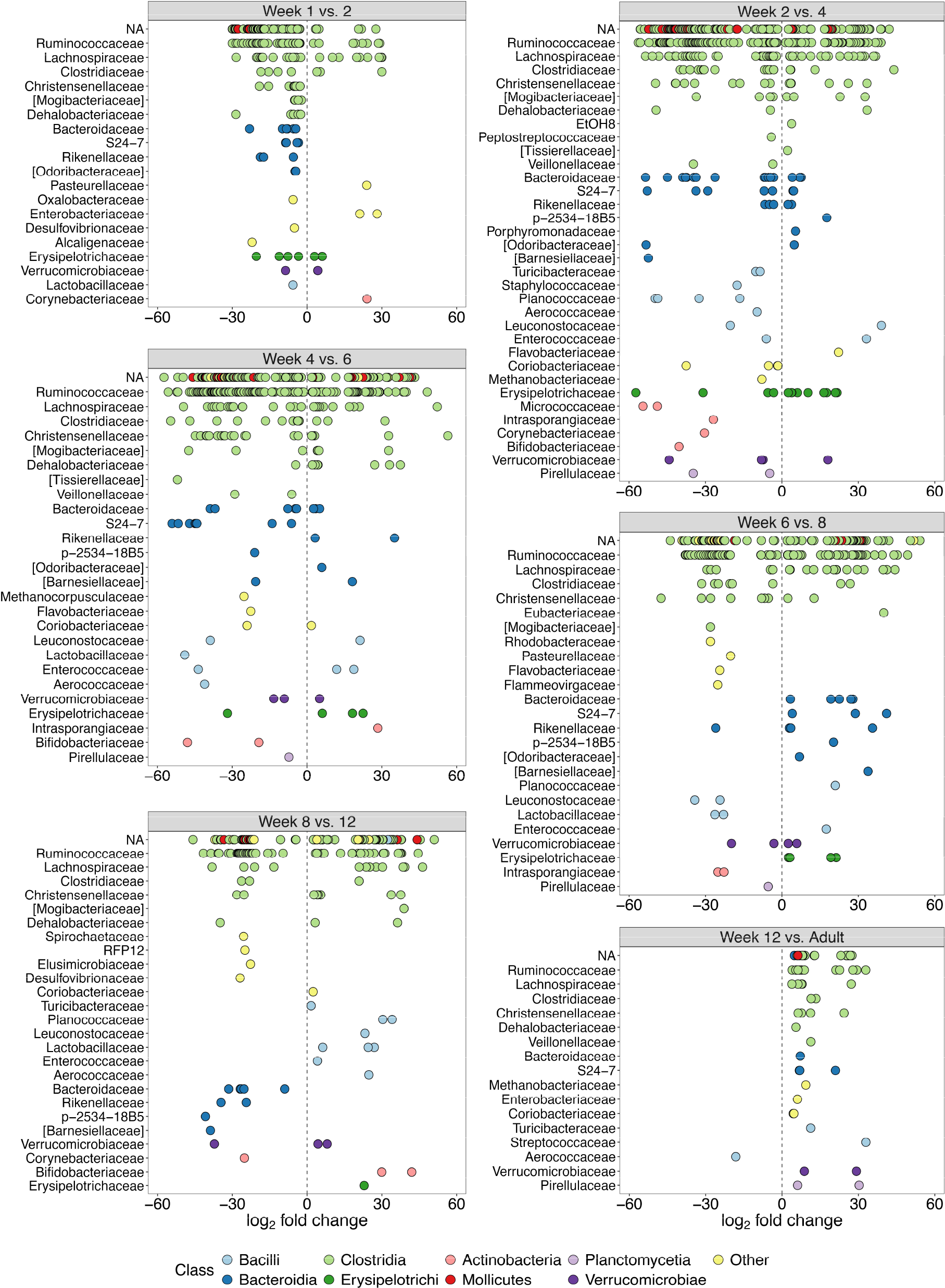
Large differences in OTU abundances between host ages closest in time. Dots show significant differentially abundant OTUs (q < 0.01) between age groups, y-axes show taxonomic families, and all OTUs have been coloured at the class level. Positive log_2_ fold changes indicate higher relative OTU abundance in the younger age group in each comparison, and negative log_2_ fold changes indicate higher abundance in the older age group. NA = OTUs without family classification.

### Associations between gut microbiota and growth of juvenile ostriches

The relationship between alpha diversity and weight varied across the different ages (lme: weight:age parameter estimate (*b*) ± *se* = −0.008 ± 0.003, F_1, 203_ = 7.49, p = 0.007). We found that at early ages there was a positive association between weight and diversity, which disappeared at later ages (Figure 8).

**Figure 8.**
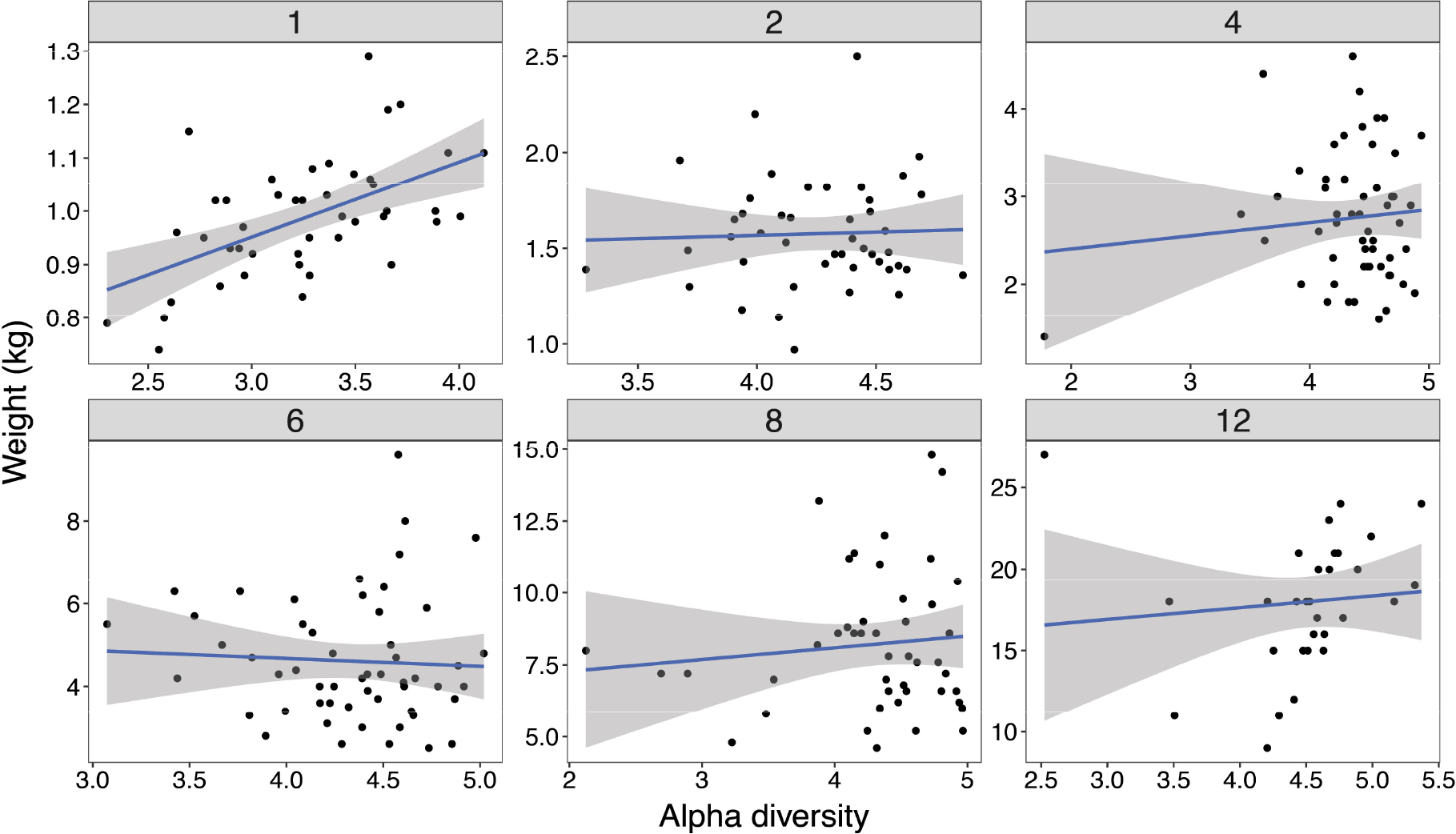
The relationship between microbial diversity and juvenile growth. Weight compared to alpha diversity (Shannon index) at each age in weeks (headers). The blue lines represent linear regression lines and the shaded areas show the 95% confidence interval. Week 1 is the only age showing significant correlation between alpha diversity and weight (r = 0.55).

Further investigation of the relationship between alpha diversity and weight using age specific correlations rendered similar results: there was a highly significant positive correlation between weight and alpha diversity at week 1 of age (r = 0.55, p = 0.0001) (Figure 8), which disappeared at subsequent ages (weeks 2, 4, 6, 8, and 12; r: −0.05-0.10, p > 0.49). Patterns of richness were very similar to those of alpha diversity with positive associations with weight shortly after hatching, but not at later ages (lme: weight:age parameter estimate (*b)* ± *se* = −1.66 ± 0.54, F_1, 203_ = 9.40, p = 0.003).

Correlation tests between log-transformed normalised OTU abundances of different taxa and weight showed that the family Bacteroidaceae was positively correlated with juvenile weight during week 1 of age (class: Bacteroidia, r = 0.47, n = 44, p = 0.001, q = 0.037) (Figure 9). In contrast, at week 2, weight was negatively correlated with the abundance of Enterobacteriaceae (class: Gammaproteobacteria, r = −0.46, n = 45, p = 0.002, q = 0.037), and at week 6, weight was negatively correlated with the abundance of two Bacilli families, Enterococcaceae (r = −0.44, n = 54, p = 0.001, q = 0.037) and Lactobacillaceae (r = −0.41, n = 54, p = 0.002, q = 0.037) (Figure 9; Table S9). Despite lower resolution at lower taxonomic levels, the genus *Bacteroides* (r = 0.47, q = 0.043) still showed strong signals of being significantly positively correlated with weight at week 1, and the genera *Enterococcus* (r = −0.43, q = 0.043) and *Lactobacillus* (r = −0.41, q = 0.043) were still significantly negatively correlated with weight at week 6 (Table S10).

**Figure 9.**
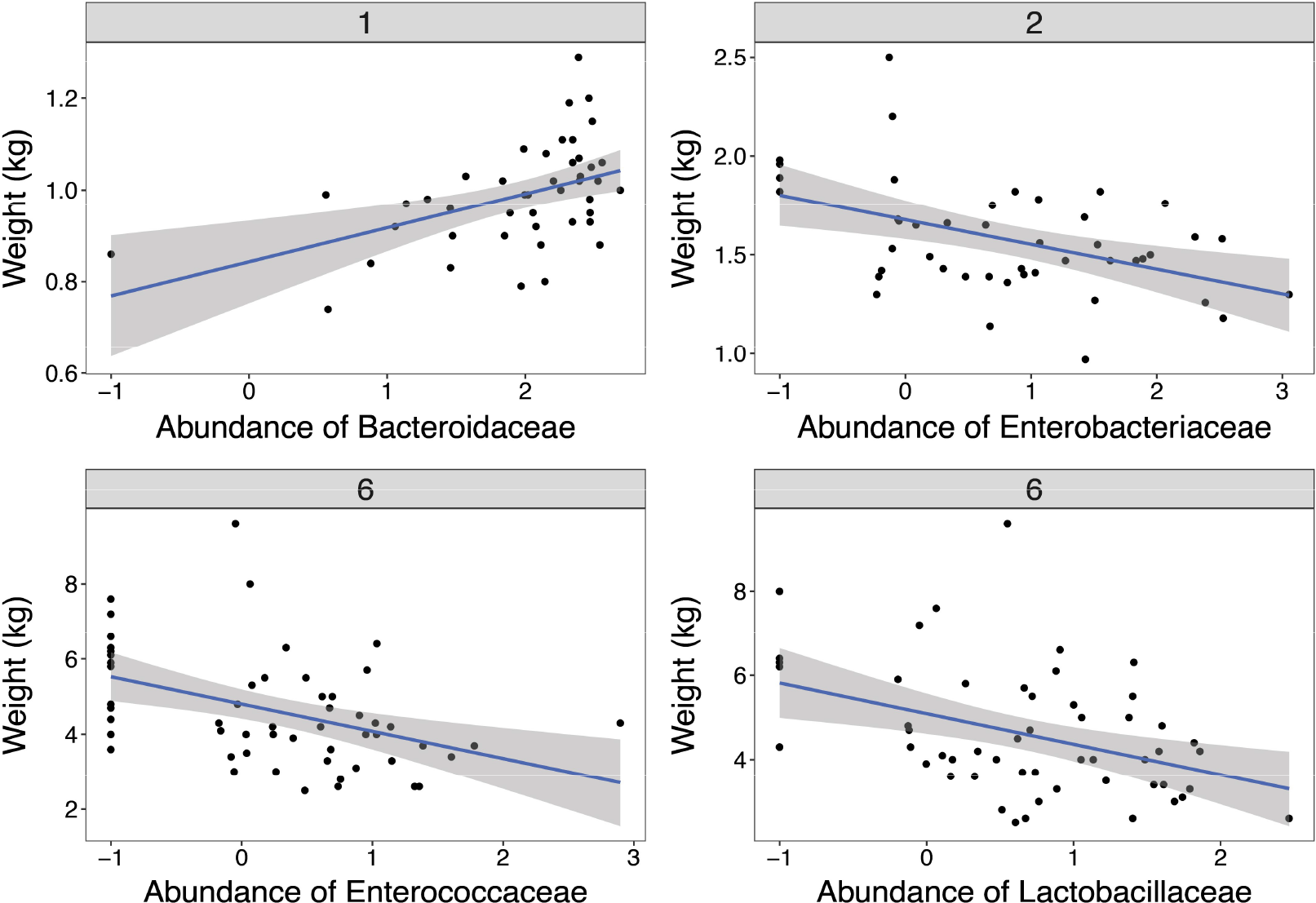
The relationship between taxon abundances and juvenile growth. Weight compared to log-transformed normalised abundances (+0.1) of the four bacterial families that showed significant (q < 0.05) correlation coefficients. The blue lines represent linear regression lines and the shaded areas show the 95% confidence interval.

## Discussion

The maturation of the gut microbiota during development is a crucial process affecting potential variation in host fitness. Studies on non-model organisms have, however, been lacking and the extent to which microbial diversity and specific bacterial taxa underlie changes in juvenile growth throughout development has been unclear. We found that bacteria colonise the gut of juvenile ostriches in a successional manner, and develops with increasing diversity and complexity as individuals age. Major compositional changes in the gut microbiota were observed, in particular from the first to the second week of life, coinciding with a dietary switch from yolk to food, and the relationship between microbiota and juvenile growth was taxon-specific and changed over time, potentially explaining some of the contradictions reflected in previous research.

During the first week of life, the gut microbiome of ostriches was highly differentiated from that of subsequent ages, with a much lower alpha diversity and a unique microbial composition dominated by Verrucomicrobiae, Clostridia, Erysipelotrichi, and Bacteroidia. Both Verrucomicrobiae and Erysipelotrichi were most abundant during this early phase, but rapidly decreased during later ages (Figure 5). At week 1 of age, ostrich juveniles are heavily dependent on yolk from their internal yolk sac for nutrition, which is high in fat and protein. After the first week, the yolk sac has largely been absorbed and they switch to external sources of food (Deeming 1999), mainly plant matter which has a high proportion of fibre. Diet has been shown to have major effects on the gut microbiome of animals (Koenig *et al.* 2011; Muegge *et al.* 2011; Jiang *et al.* 2017), so it is likely that the dietary switch during this time has a direct impact on the differences we see in the gut community of 1- and 2-week-old chicks. Partly because of this switch from yolk to external food sources, it is generally recognized that the early post-hatch period is a critical stage for the growth and health of poultry (Gilbert *et al.* 2010; Cheled-Shoval *et al.* 2011; Pan & Yu 2014). It has been demonstrated in chickens, for example, that the digestive organs of newly hatched juveniles undergo both large anatomical and physiological changes to accommodate this dietary transition (Uni *et al.* 1999).

One of the most striking changes in the development of the gut microbiota was exhibited by the Verrucomicrobiae. This class consists of only a single species in our data, specifically *Akkermansia muciniphila*, which dominates the gut of 1-week-old ostriches (36.1% in total), while being almost non-existent in the adult individuals (0.09%; Figure 5). *A. muciniphila* is a mucin degrader found in a wide variety of animal species (Belzer & de Vos 2012), and has been well studied in the gut microbiota of mice and humans for its anti-inflammatory effects and negative correlation with obesity, diabetes, and inflammatory gut diseases (Everard *et al.* 2013; Schneeberger *et al.* 2015; Derrien *et al.* 2017). For example, a study by Caesar *et al.* on mice showed that *A. muciniphila* was positively associated with a diet rich in polyunsaturated fat, and faecal transplants from these mice corresponded to reduced inflammation levels in recipient mice regardless of their diet (Caesar *et al.* 2015). This taxon is also hypothesized to protect the gut from pathogens through competition (Belzer & de Vos 2012), which could be an important mechanism in the guts of young chicks at such a sensitive stage of development. We are not aware of any study linking *A. muciniphila* with a diet rich in yolk, but the high prevalence in 1-week-old ostrich juveniles and the subsequent rapid decline seems to suggest a possible association. Further research is needed to establish this link and whether it has similar beneficial effects in ostriches as those documented in mice and humans.

The adult ostrich faecal microbiome is heavily dominated by Clostridia, primarily the families Ruminococcaceae, Lachnospiraceae, and Clostridiaceae, with a minor prevalence of Bacteroidia and other classes (Figure 4). This taxonomic composition broadly agrees with the faecal bacterial composition of other hindgut fermenters (O’ Donnell *et al.* 2017), and the caecal microbiome of the chicken (Gong *et al.* 2007; Ballou *et al.* 2016), turkey (Scupham 2007), and ostrich as reported in a previous study (Matsui *et al.* 2010). It differs, however, from the study by Bennett *et al.* (2013) who evaluated the caecal microbiota of another ratite, the emu (Dromaius novaehollandiae), where Clostridia only comprised a minor proportion of sequences. In our previous work on the microbiota of gastrointestinal regions from juvenile ostriches, we showed that the caecum harbours a differentiated bacterial community compared to that of faeces (Videvall *et al.* 2017a; b), so any differences between faecal and caecal microbiota across host species will be very difficult to interpret. Further studies characterising ratite gut microbiota are needed in order to establish whether the ostrich microbial composition is similar to that of related species.

Previous research has found contrasting effects of microbial diversity on host development and growth. Our results showed that the developmental stage at which you examine the diversity of gut microbiota is crucial for understanding this relationship. Overall there was a small negative association between growth and alpha diversity, which is in line with multiple studies showing that juveniles have higher growth when the diversity of gut bacteria is reduced with antibiotics (Gaskins *et al.* 2002; Dibner & Richards 2005). However, our results demonstrate a more complicated picture, with bacterial diversity having strong positive effects on growth just after hatching when the gut microbiota is relatively simple. Analyses of bacterial abundances could pinpoint, however, specific taxa that were associated with growth at certain ages. As an example, individuals with higher abundances of *Lactobacillus* had reduced weight, a finding which agrees with numerous studies of chickens, where it has been well-documented that species of *Lactobacillus* negatively affect growth via intestinal bile salt hydrolase activity (Engberg *et al.* 2000; Guban *et al.* 2006; Danzeisen *et al.* 2011; Lin *et al.* 2013; Crisol-Martínez *et al.* 2017). A meta-analysis of multiple *Lactobacillus* species have demonstrated, however, that some are associated with an increase, and some with a decrease in weight across different animals (Million *et al.* 2012; see also Angelakis & Raoult 2010). Further studies are clearly needed to isolate and validate specific bacterial strains related to juvenile growth. Although our results are correlational and so we cannot infer causality, it highlights the importance of examining specific taxa, microbial diversity, and host traits, such as growth, in concert at specific developmental windows, rather than characterising broad taxonomy profiles across periods that encompass different developmental stages.

In summary, this study describes the successional development of the gut microbiota in juvenile ostriches during their first three months of life. We have showed a gradual maturation of microbial diversity and richness, multiple microbial colonisation and extinction events, and major taxonomic shifts co-occurring with a dietary switch. There were different associations between the diversity of gut microbiota and juvenile growth, and we identified specific bacterial groups contributing to this relationship. Our study indicates that colonisation and extinction processes in the gut occurs in succession, and together with the fact that different bacteria were found to have positive and negative effects on host fitness, indicates that changes in the community over time may be the outcome of an interplay between bacterial and host interests. These results have implications for our understanding of gut microbiome development of hosts and how it influences fitness-related traits such as growth. Future research is needed to investigate potential sources of microbial recruitment, and the causal mechanisms determining microbiota abundance at different stages of development.

## Acknowledgements

We are grateful to all staff at the Oudtshoorn Research Farm, Western Cape Government, for assisting with sample collection. Funding was provided by the Helge Ax:son Johnson Foundation, the Längmanska Cultural Foundation, the Lund Animal Protection Foundation, the Lars Hierta Memorial Foundation, and the Royal Physiographic Society of Lund to E.V., by a Wallenberg Academy fellowship and a Swedish Research Council grant to C.K.C., and by the Western Cape Government through the use of their facilities and animals.

## Author contributions

E.V. and C.K.C. planned and designed the study. S.C. provided animal facilities. A.E. supervised the experimental part of the study. N.S., A.E., C.K.C., and E.V. performed the sampling and cared for the animals. A.O. advised on sampling procedure. M.S. supervised the laboratorial part of the study, and together with H.M.B. prepared the samples for sequencing. E.V. performed the bioinformatic and statistical analyses. C.K.C., S.J.S., R.K., and O.H. provided advice on analyses and the interpretation of results. E.V. and C.K.C. wrote the paper with input from all authors.

